# Synthesis and application of a photocaged L-lactate

**DOI:** 10.1101/2024.01.30.577898

**Authors:** Ikumi Miyazaki, Kelvin K. Tsao, Yuki Kamijo, Yusuke Nasu, Takuya Terai, Robert E. Campbell

## Abstract

l-Lactate, once considered a metabolic waste product of glycolysis, is now recognized as a vitally important metabolite and signaling molecule in multiple biological pathways. However, exploring l-lactate’s emerging intra- and extra-cellular roles is hindered by a lack of tools to locally perturb l-lactate concentration intracellularly and extracellularly. Photocaged compounds are a powerful way to introduce bioactive molecules with spatial and temporal precision using illumination. Here, we report the development of a photocaged derivative of l-lactate, 4-methoxy-7-nitroindolinyl l-lactate (MNI-l-lac), that releases l-lactate upon UV illumination. We validated MNI-l-lac in cell culture by demonstrating that the photorelease of l-lactate elicits a response from genetically encoded extra- and intracellular l-lactate biosensors. These results indicate that MNI-l-lac may be useful for perturbing the concentration of endogenous l-lactate in order to investigate l-lactate’s roles in metabolism and signaling pathways.

## Introduction

l-Lactate is an abundant metabolite that is produced from pyruvate, the end-product of glycolysis, through the action of lactate dehydrogenase. While lactate has traditionally been viewed as a waste byproduct of glucose metabolism,^[1]^ recent studies suggest it has diverse roles as both an energy source and a signaling molecule in the nervous system,^[2]^ tumor microenvironment,^[3,4]^ and immune system.^[5,6]^ These emerging roles have implications for physiological and pathological processes that span intracellular,^[7]^ intercellular,^[2,8]^ and interorgan^[9,10]^ scale environments.

The increased recognition of the various biological roles of lactate has highlighted the need for improved tools to study its roles through both its measurement and perturbation in tissues. We and others have recently reported a variety of genetically encoded biosensors for the measurement and imaging of lactate in tissues.^[11–24]^ Applications of these measurement tools would be complemented by a tool for perturbing lactate concentrations. A molecule that enables the photorelease of lactate (that is, a ‘caged’ lactate) could be a particularly powerful tool for the spatiotemporal manipulation of lactate concentration in tissues. Photocaged compounds have been used as perturbation tools in a wide range of applications.^[25–28]^ Since the original reports of *o*-nitrobenzyl-caged ATP and cAMP in the late 1970s,^[29,30]^ the demand for alternative or improved caging groups has led to many new types of photocages, including ones that are coumarin-based,^[31]^ cyanine-based,^[32]^ and *o*-nitro-2-phenethyl-based.^[33,34]^

One of the most common applications of caged compounds is the study of neuronal processes by one- or two-photon uncaging of 4-methoxy-7-nitroindonyl l-glutamate (MNI-Glu), which was independently reported by Matsuzaki *et al*.^[35]^ and Canepari *et al*.^[36]^ in 2001. MNI-Glu is the most widely used type of caged glutamate due to its high stability at physiological pH, resistance to hydrolysis, and solubility in physiological buffers.^[37]^ In addition, MNI-Glu’s uncaging wavelength in the near UV is spectrally compatible with other commonly used chromophores, such as green fluorescent protein (GFP), enabling the combined use of photocaged glutamate and GFP or GFP-based biosensors.^[35]^ Other MNI-caged compounds have also been developed, including MNI-D-aspartate^[38]^ and MNI-auxins.^[39]^

In an effort to facilitate studies of lactate’s role as a metabolite and a signaling molecule, and potentially help resolve ongoing controversies such as the astrocyte to neuron lactate shuttle (ANLS) hypothesis,^[2]^ we undertook the development of an MNI-caged lactate (MNI-l-lac). In this work, we report the synthesis and characterization of MNI-l-lac and validation of lactate release in cell culture.

## Results and Discussion

### MNI-l-lac synthesis

To design a photocaged lactate, we considered two key factors: the choice of the photocaging group and the position of attachment to lactate. Based on the success and widespread application of MNI-Glu,^[35,36]^ we chose to pursue an analogous design in which the MNI photocaging group would be linked to the carbonyl carbon of lactate.

To synthesize MNI-l-lac, we expected to be able to employ a synthetic strategy analogous to the one commonly reported for MNI-Glu.^[35,36]^ Specifically, we planned to couple 4-methoxyindoline to lactate with an amide bond-forming reaction. We would then perform a nitration reaction to introduce the 7-nitro group at the desired position, *para* to the 4-methoxy group. Following this plan, we succeeded in coupling 4-methoxyindoline with lactate. However, when we attempted the nitration reaction, we found that the nitration occurred preferentially on the hydroxyl group of lactate rather than at the desired position on the 4-methoxyindoline. To circumvent this problem, we tried nitrating 4-methoxyindoline first before coupling the nitrated product to free lactate. However, we were unable to obtain the desired MNI-l-lactate, possibly due to the electron-withdrawing nature of the nitro group attenuating the indoline’s nucleophilicity.

Ultimately, we obtained the desired product by first protecting the hydroxyl group of ethyl lactate with *tert*-butyldimethylsilyl (TBDMS) to give compound **2** (**Figure 1a**). 4-methoxyindoline (compound **3**) was prepared by reducing commercially available 4-methoxyindole with sodium cyanoborohydride in acetic acid. Compound **3** was then successfully coupled with compound **2** using HATU/EDC coupling conditions to give compound **4**. Compound **4** was nitrated using silver nitrate and acetyl chloride, which gave a mixture of the *ortho*- and *para*-nitro-substituted regioisomers (compounds **5** and **6**, respectively). These regioisomers were separated by flash chromatography using silica gel to yield the *ortho*-isomer as an oil (compound **5**) and the *para*-isomer as a yellow solid (compound **6**). The desired regioisomer, compound **6**, was deprotected with TBAF/AcOH to afford MNI-l-lac (compound **7**) as a yellow solid. The deprotection was performed under acidic conditions because standard basic conditions resulted in cleavage of the amide bond, yielding free 7-nitroindoline as the major product. This lability under basic conditions suggests that MNI-l-lac is susceptible to amide bond cleavage, which may explain the poor yield for the coupling reaction between unprotected lactate and indoline in the initial (and unsuccessful) synthetic strategy.

**Figure 1.**
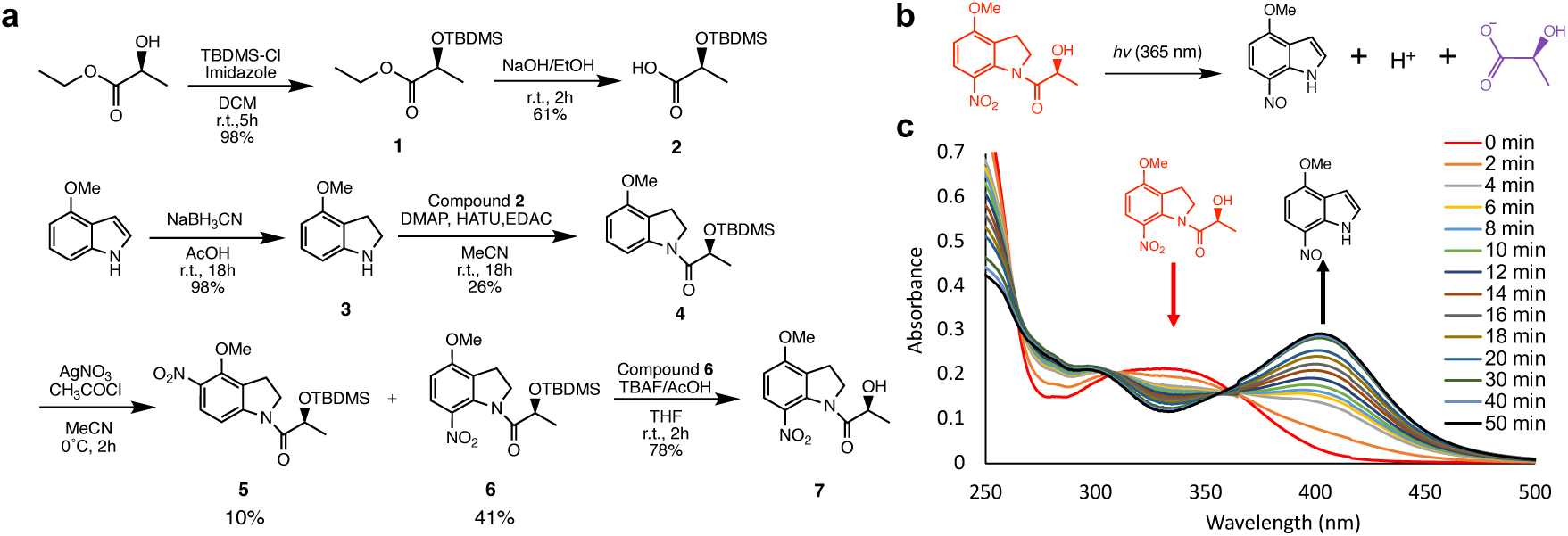
Design and validation of MNI-l-lac. (**a**) Synthetic scheme of MNI-l-lac. (**b**) Schematic representation of photouncaging of MNI-l-lac to release lactate plus a proton. (**c**) Absorbance spectra were recorded during the progressive photolysis of MNI-l-lac (40 μM solution in lactate (−) buffer (30 mM MOPS, 100 mM KCl, 1 mM CaCl2, pH 7.2) containing 10% DMSO).

### MNI-l-lac characterization

To investigate whether the expected uncaging reaction^[40]^ occurred upon UV illumination of MNI-l-lac, we measured the UV-Vis absorption spectra of the compound exposed to 365 nm UV light for increasing durations (**Figure 1b**). The progressive photolysis of MNI-l-lac in aqueous neutral buffer results in a bathochromic shift in the absorption maximum, with distinct isosbestic points (**Figure 1c**). Based on this data, we estimate that the conversion is 90% complete within 10 min illumination with 365 nm light (810 μW/cm^2^). We also observed the color of the MNI-l-lac solution turn from clear to yellow upon UV illumination, consistent with the bathochromic shift in the absorption spectra and the conversion of the starting MNI-l-lac (λ_max_ = 330 nm) into the 4-methoxynitrosoindole byproduct (λ_max_ = 403 nm).^[41–43]^

To confirm that lactate was released upon UV illumination of MNI-l-lac, we used a commercial l-lactate assay kit to quantify the amount of lactate released in both buffered and unbuffered aqueous solutions at two concentrations (**Supplementary Figure 1** and **Supplementary Table 1**). The results indicated that lactate was released upon illumination to over 86 ± 8% (mean ± SD) of the theoretically expected concentration. The difference between the uncaging efficiency in buffered versus unbuffered solution was insignificant (one-way ANOVA, p = 0.858).

### *In vitro* characterization of MNI-l-lac using a lactate biosensor

We envisioned that our previously reported genetically encoded lactate biosensors, such as eLACCO1 (**Table 1**), would provide an effective means of characterizing and validating the utility of MNI-l-lac in the context of cultured cells.^[15]^ Before proceeding with cell-based experiments, we first needed to confirm that MNI-l-lac could be uncaged in the presence of the purified eLACCO1 biosensor (**Figure 2a**).^[15]^ These tests are also necessary to validate that eLACCO1 only responds to the resulting uncaged lactate and establish any effects that could arise from excess MNI-l-lac.

**Figure 2.**
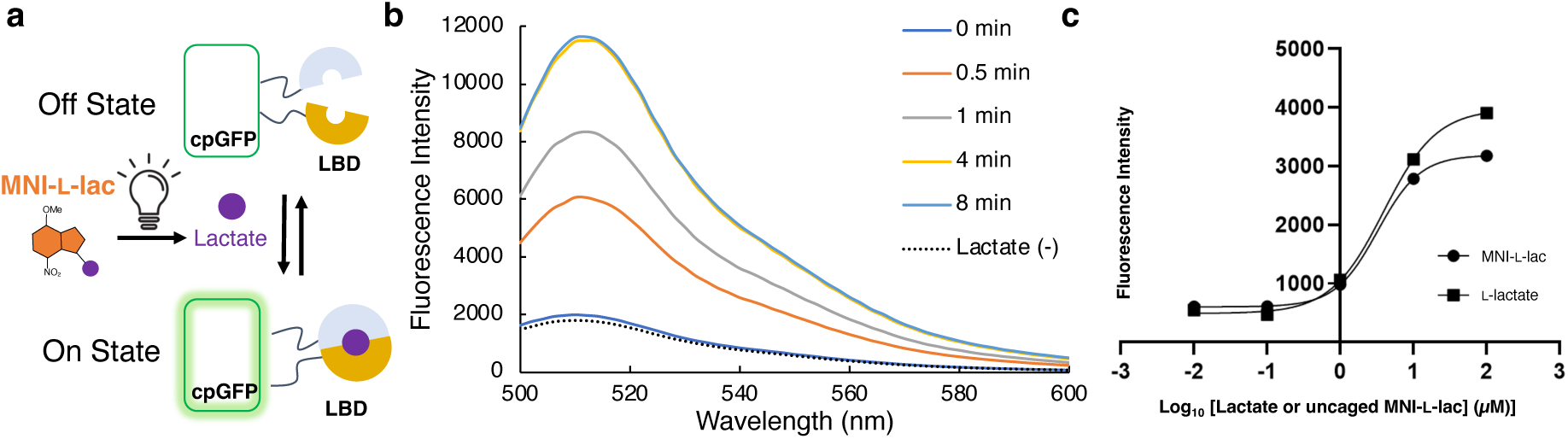
*In vitro* characterization of MNI-l-lac. (**a**) Schematic representation of the eLACCO1 biosensor’s fluorescence response. eLACCO1 consists of cpGFP + lactate binding domain (LBD). (**b**) eLACCO1 fluorescence response with MNI-l-lac (40 µM) with different UV illumination times (average of *n* = 3). (**c**) Overlay of lactate (squares) and uncaged MNI-l-lac (circles) titrations using the eLACCO1 fluorescence biosensor (*n* = 3). MNI-l-lac solutions in varying concentrations were illuminated for 10 minutes.

**Table 1.**
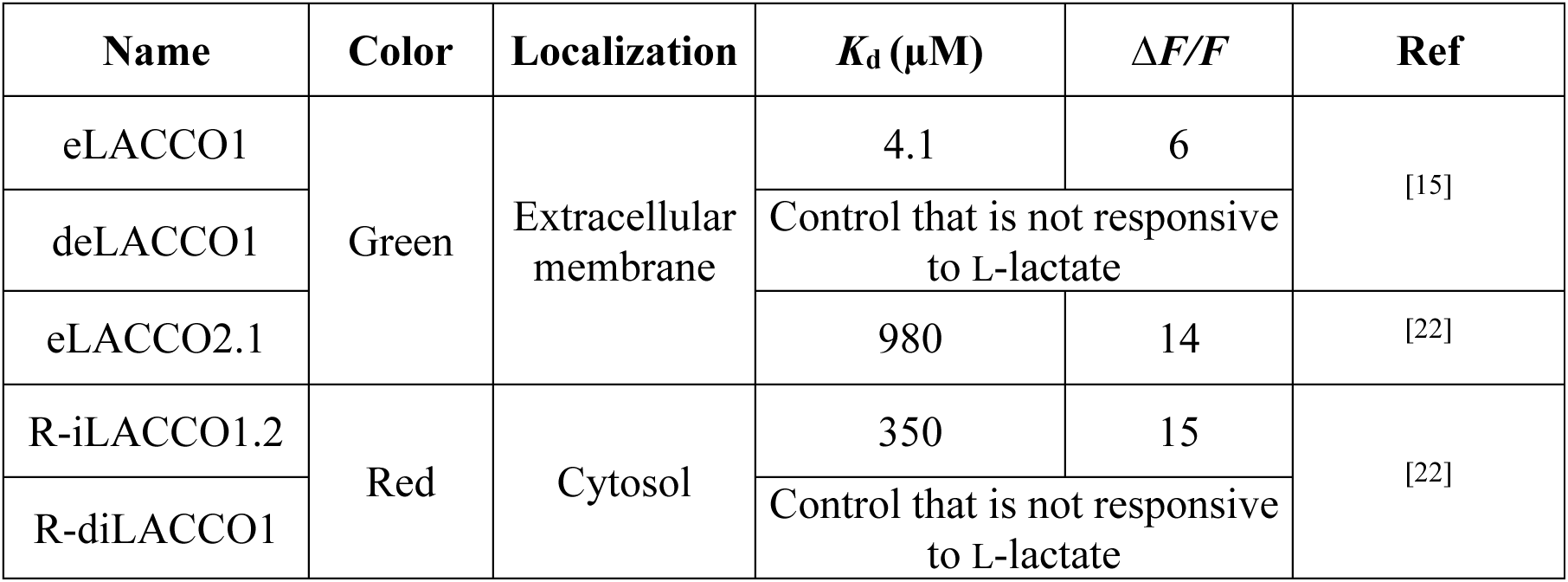
Lactate biosensors used in this work.

| Name | Color | Localization | $K_d$ (μM) | $\Delta F/F$ | Ref |
| --- | --- | --- | --- | --- | --- |
| eLACCO1 | Green | Extracellular membrane | 4.1 | 6 | [15] |
| deLACCO1 |  |  | Control that is not responsive to L-lactate |  |  |
| eLACCO2.1 |  |  | 980 | 14 | [22] |
| R-iLACCO1.2 | Red | Cytosol | 350 | 15 | [22] |
| R-diLACCO1 |  |  | Control that is not responsive to L-lactate |  |  |

To determine if the MNI-modified lactate was indeed “caged” with respect to binding to the eLACCO1 biosensor, we performed a fluorescence assay using purified eLACCO1 protein along with MNI-l-lac exposed to varying durations of UV light. The result revealed that the fluorescence intensity of eLACCO1 with MNI-l-lac (*t* = 0 min; no light illumination) and the lactate (−) control condition (lactate-free state) was very similar (**Figure 2b**). This indicated that MNI-l-lac itself did not induce a meaningful fluorescence response for eLACCO1. Additionally, as the illumination time of MNI-l-lac increased, so too did the fluorescence response of eLACCO1. This increase in fluorescence indicates eLACCO1’s response to the release of free lactate from MNI-l-lac upon light illumination. The fluorescence response of the eLACCO1 biosensor reached its maximum after 8 minutes of UV illumination. Accordingly, for all later experiments, we used 10 minutes of illumination time to ensure near-complete uncaging of MNI-l-lac.

To further investigate whether lactate released from MNI-l-lac is indeed l-lactate (as opposed to D-lactate), we performed a titration experiment against eLACCO1 to determine the apparent dissociation constant (*K*_d_) value. If MNI-l-lac releases l-lactate, we expect identical *K*_d_ values and maximum fluorescence responses (τι*F*/*F*) for completely uncaged MNI-l-lac (i.e., illuminated for 10 min with 365 nm light) and native l-lactate. The fluorescence intensity of lactate titration using sodium l-lactate and uncaged MNI-l-lac were plotted against log [l-lactate or uncaged MNI-l-lac] (µM), which was fitted to the Hill Equation to find the *K*_d_ values (**Figure 2c**). The apparent *K*_d_ values were calculated to be 4.0 µM and 3.3 µM for l-lactate (previously reported values were 4.1 µM for l-lactate, 120 μM for D-lactate) and uncaged MNI-l-lac, respectively.^[15]^ Similarly, the *ΔF*/*F*_0_ of eLACCO1 at the maximum concentration of 100 µM was calculated to be 7.2 for l-lactate and 5.7 for uncaged MNI-l-lac, which were close to the literature value of 6.^[15]^ There appeared to be a substantial divergence in the fluorescence response with standard l-lactate versus uncaged MNI-l-lac at concentrations above 10 µM, which may be due to fluorescence attenuation by an inner filter effect. This is suggested by **Figure 1c**, where the byproduct 4-methoxynitrosoindole has some absorbance over 450 nm that partly overlaps the excitation of eLACCO1. Overall, these results are consistent with the conclusion that MNI-l-lac can be used to release free l-lactate at concentrations up to approximately 100 µM.

### Cell-based characterization of MNI-l-lac using lactate biosensors

Genetically encoded fluorescent biosensors enable precise and dynamic imaging of lactate in either the extracellular or intracellular environments, allowing for detailed spatial and temporal analysis. We have previously reported the green-fluorescent eLACCO series of biosensors for extracellular lactate,^[15,22]^ and the red-fluorescent R-iLACCO series for intracellular lactate (**Table 1**).^[22]^ Our *in vitro* results had confirmed that illumination of MNI-l-lac releases lactate, which, in turn, can bind to and cause the high-affinity eLACCO1 biosensor to respond.

We next aimed to determine if MNI-l-lac could be used to perturb the lactate concentration in the environment of cultured mammalian cells. To determine if uncaging of MNI-l-lac could perturb the extracellular lactate concentration, we prepared HeLa cells expressing the lactate biosensor eLACCO2.1 or the non-lactate responsive deLACCO1 control on the cell surface. Relative to eLACCO1, eLACCO2.1 has a higher *K*_d_ which is within the range of relevant extracellular lactate concentrations (**Table 1**). We performed fluorescence imaging to determine the response of deLACCO1 and eLACCO2.1 on the surface of HeLa cells to uncaging of MNI-l-lac (1 mM, 1 s, 405 nm, 2.4 mW/cm^2^). The deLACCO1 variant is not responsive to lactate but retains the pH sensitivity of eLACCO2.1.^[15]^ Controlling for pH changes in this experiment is essential due to the fact that uncaging of MNI-l-lac releases lactate plus a proton (**Figure 1b**). In addition, lactate is transported across the mammalian cell membrane by proton-dependent monocarboxylate transporters (MCTs), such that lactate import is necessarily accompanied by proton import and a decrease in cytosolic pH.^[44,45]^

When MNI-l-lac was uncaged in the presence of HeLa cells, we observed distinctly different fluorescent responses for cells expressing eLACCO2.1 versus deLACCO1 (**Figure 3a**). For cells expressing the deLACCO1 control biosensor, uncaging produced a rapid 37 ± 1% (mean ± SEM, *n* = 10) decrease in fluorescence followed by a recovery to baseline within about one minute. For cells expressing the eLACCO2.1 biosensor, uncaging produced a smaller 10 ± 3% (mean ± SEM, *n* = 9) decrease in fluorescence followed by an increase to a value that was 19 ± 4% (mean ± SEM, *n* = 9) greater than the baseline. We rationalize these responses as follows. The first sudden decrease in Δ*F*/*F* immediately after photoactivation is likely due to the release of protons from the uncaging reaction of MNI-l-lac, causing a fluorescence decrease in these pH sensitive biosensors.^[15]^ Second, the transient increase in fluorescence observed with eLACCO2.1 but not with deLACCO1, suggests that the local concentration of lactate has been substantially increased due to the released lactate from MNI-l-lac. The later slow decrease in fluorescence may be due to the diffusion of lactate over time.

**Figure 3.**
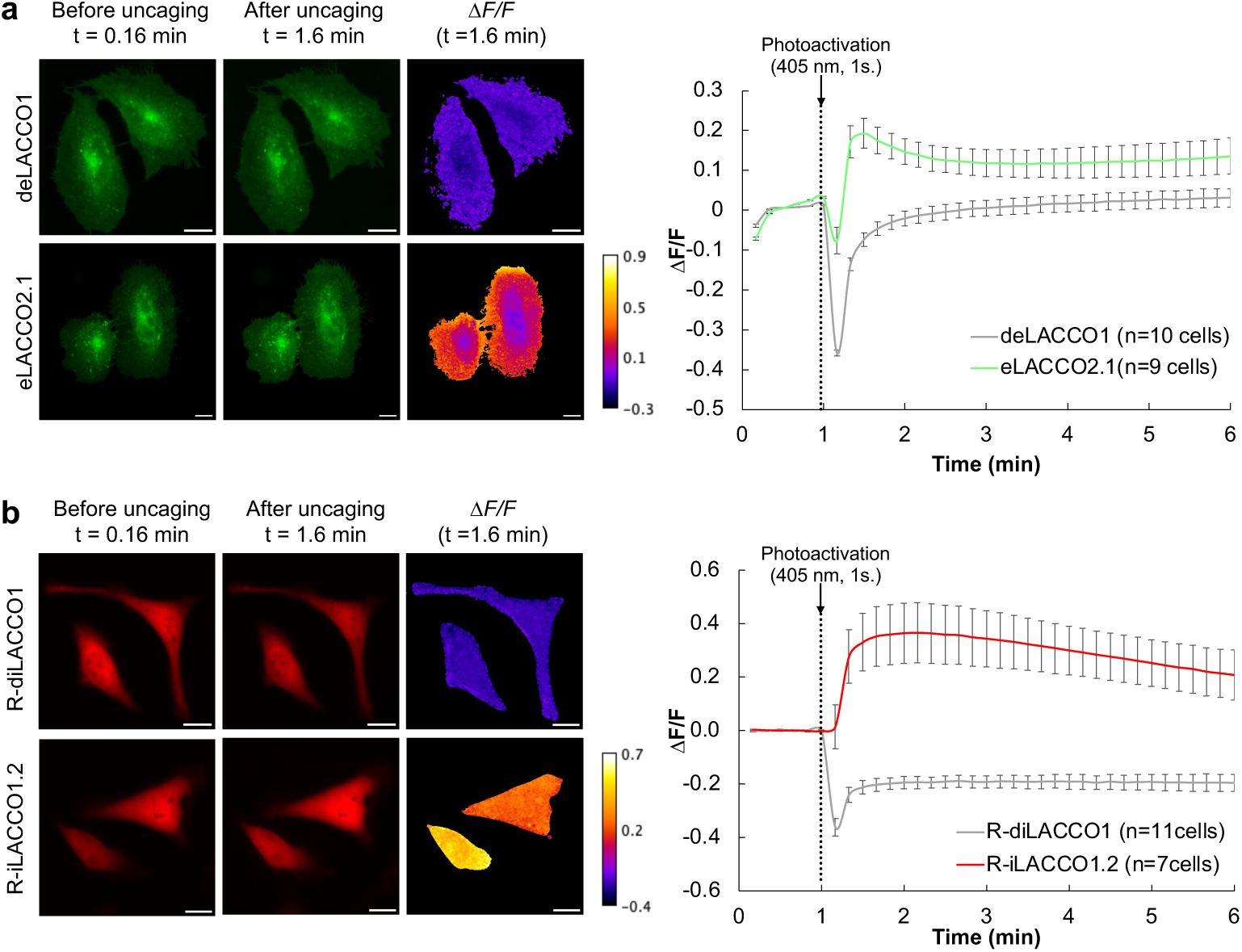
Cell-based characterization of MNI-l-lac. (**a**) deLACCO1 and eLACCO2.1 expressed on the cell surface of HeLa cells before and after MNI-l-lac photoactivation at 1 minute (405 nm, 1 s, 2.4 mW/cm^2^). *n* = 10 and 9 cells for deLACCO1 and eLACCO2.1, respectively (mean ± SEM). (**b**) R-diLACCO1 and R-iLACCO1.2 expressed in HeLa cells before and after MNI-l-lac photoactivation at 1 minute (405 nm, 1 s, 2.4 mW/cm^2^), in the absence of AR-C155858. *n* = 11 and 7 cells for R-diLACCO1 and R-iLACCO1.2, respectively (mean ± SEM). Photoactivation was performed over the entire field of view. Images in the third column of (**a**) and (**b**) are pseudocolored to demonstrate the change in fluorescence before photouncaging (*F0*, t = 0.16 min) and after photouncaging (*F*, t = 1.6 min). The values shown in these images were calculated as (*F-F0*)/*F0*.

As an additional control, the imaging experiment was performed in the absence of MNI-l-lac. The first decrease of Δ*F/F* by 2 ± 1% (mean ± SEM, *n* = 13) for deLACCO1 and 14 ± 0.2% (mean ± SEM, *n* = 12) for eLACCO2.1 was still evident after photoactivation but less than in the experiment with MNI-l-lac (**Supplementary Figure 2**). One plausible explanation for this decrease is the pH dependent quenching of the GFP in biosensors, causing a transient reduction in fluorescence. Furthermore, while we did not observe a transient increase in fluorescence for HeLa cells expressing eLACCO2.1, there was a gradual increase in the value of 18 ± 2% (mean ± SEM, *n* = 12) over time post-photoactivation. This is attributed to the efflux of endogenous l-lactate from cells to extracellular space as cells continually produce l-lactate through glycolysis over the course of measurement. The reasoning is also supported by a small but steady increase of eLACCO2.1 (not observed in deLACCO1) fluorescence even before photoirradiation (**Figure 3a**, 0-1 min). In summary, these results support the conclusion that uncaging of MNI-l-lac releases lactate at a sufficient concentration to induce the fluorescence response of eLACCO2.1.

Next, we characterized the ability to visualize the increase in intracellular lactate concentration upon uncaging MNI-l-lac with HeLa cells expressing R-iLACCO1.2 and R-diLACCO1. Contrary to the eLACCO cell-based characterization, we imaged HeLa cells expressing R-iLACCO1.2 under glucose-starved conditions to reduce the intracellular lactate concentration and to better observe the change upon uncaging MNI-l-lac (5 mM). In addition, R-iLACCO imaging was performed with and without AR-C155858,^[46]^ an inhibitor of the MCT1 and MCT2 lactate transporters. With no inhibitor, R-iLACCO1.2 exhibited a 21 ± 5% (mean ± SEM, *n* =12) increase in its fluorescence response to lactate released due to the illumination of MNI-l-lac (**Figure 3b**). However, in the presence of MCT inhibitor, R-iLACCO1.2 no longer responded to the uncaging event (**Supplementary Figure 3**). The result can be explained by assuming that MNI-l-lac has poor cell membrane permeability and remains primarily outside the cell until it is uncaged. When MNI-l-lac uncages in the absence of AR-C155858, the photoreleased lactate would be shuttled into the cells through the MCT transporters. In the absence of MCT inhibitor condition (**Figure 3b**), both R-iLACCO1.2 and R-diLACCO1 resulted in a sudden decrease in fluorescence of 5 ± 3% (mean ± SEM, *n* = 12) and 38 ± 1% decrease (mean ± SEM, *n* = 20) respectively, which are consistent with what was seen when using eLACCO. This sudden decrease is, again, likely due to the influx of protons into the cell, which disrupts the fluorescence of the pH-sensitive R-iLACCO despite the uncaging event occurring outside the cell. Additionally, the persistent negative fluorescence change observed in the lactate unresponsive R-diLACCO1 is likely due to the intracellular pH reaching a new equilibrium, stabilizing the fluorescence at a lower level than before the photoactivation of MNI-l-lac. We see this phenomenon once more when MNI-l-lac is uncaged but in the presence of MCT inhibitor (**Supplementary Figure 3**) but in both sensors instead, since lactate is unable to enter the cells.

When the additional control experiment was performed in the absence of MNI-l-lac, no substantial change in fluorescence was observed for HeLa cells expressing R-diLACCO1 control variant (**Supplementary Figure 4**). As expected, when there is no MNI-l-lac present, there should be no transient pH decrease due to the release of a proton from the uncaging reaction. However, we noticed a persistent decrease in fluorescence for the lactate-responsive R-iLACCO1.2 after photoactivation, likely caused by photobleaching of the R-iLACCO1.2’s RFP. Since this effect was not observed with the dead control, R-diLACCO1 may not serve as an effective comparison in this context. Regardless, R-iLACCO1.2 effectively responded to the uncaging of MNI-l-lac, validating its intracellular applicability within the cellular environment.

## Conclusion

We have synthesized, characterized, and validated the first photocaged lactate molecule, 4-methoxy-7-nitroindolinyl-l-lactate (MNI-l-lac). The progressive photolysis studies, along with the quantification of released l-lactate using a commercial assay, confirmed the ability of MNI-l-lac to release free l-lactate only after light illumination. The lactate biosensors were then used to validate the release of lactate *in vitro*. The result demonstrated a correlation between the fluorescence response and the uncaging of MNI-l-lac, validating its compatibility with the visualization tool. Furthermore, we demonstrated the combined use of MNI-l-lac to locally perturb the lactate concentration in cultured cells and visualize the change of lactate concentration with the lactate biosensors. The results from utilizing both intra- and extra-cellular lactate biosensors demonstrate that MNI-l-lac is likely uncaged outside the cells and then shuttled inside the cells. MNI-l-lac is a powerful new tool that will further enable researchers to explore the spatial and temporal dynamics of lactate as a signaling molecule and metabolite in live tissues.

## Methods

### General Methods and Materials

All chemicals and solvents were purchased from Tokyo Chemical Industries, Wako Pure Chemical, and Aldrich Chemical Co. Nuclear magnetic resonance (NMR) spectra were recorded on a JEOL JNM-ECS400 or JEOL JNM-ECZ500R/M3 instrument. Mass spectra were measured with a Bruker Compact System. Products were purified by flash column chromatography using Isolera-1SW (Biotage) and Biotage Sfär Silica HC D column. Absorbance spectra were measured using the Shimadzu UV1800 spectrometer. Fluorescence emission spectra were recorded on a Spark plate reader (Tecan).

### Synthesis of MNI-l-lac

#### TBDMS-l-lactate (compound 2)

TBDMS-l-lactate synthesis was adapted from reported procedures^[47,48]^. To a 0 °C solution of ethyl-l-lactate (2 g, 17 mmol) in DCM (20 mL), *tert*-butyldimethylsilyl chloride (TBDMS-Cl) (2.7 g, 18 mmol) and imidazole (1.4 g, 21 mmol) were added. The mixture was brought to r.t., stirred for 2 h before diluting with H_2_O (60 mL), and extracted with DCM (3 × 20 mL). The combined organic layers were washed with ice-cold 5% HCl solution, followed by washing with brine. After, the organic layers were dried over anhydrous Na_2_SO_4_, filtered, and concentrated under reduced pressure. The crude product (compound **1**) was dissolved in EtOH (10 mL) and added slowly to an NaOH solution (10 mL, 8 eq). The reaction mixture was stirred for 2 h at room temperature, then acidified to pH 2.0 with ice-cold 1 M HCl solution and extracted with EtOAc. The combined organic layers were washed with brine, dried over anhydrous Na_2_SO_4_, filtered, and concentrated under reduced pressure to afford compound **2** as an oil in a 60% yield over two steps (0.6 g, 1.83 mmol). The product (compound **2**) was used without further purification.

^1^H-NMR (400 MHz, CDCl_3_) δ 4.36 (t, J = 6.6 Hz, 1H), 1.45 (d, J = 6.4 Hz, 3H), 0.92 (s, 9H), 0.14 (s, 6H)

#### 4-methoxyindoline (compound 3)

4-Methoxyindoline (compound **3**) was synthesized according to a reported experimental procedure:^[35]^ To a solution of 4-methoxyindole (512 mg, 3.5 mmol) in acetic acid (7.5 mL), sodium cyanoborohydride (219 mg, 3.5 mmol) in DCM (10 mL) was added slowly over 15 min. The mixture was stirred at room temperature overnight. The reaction mixture was then adjusted to pH 14 with NaOH solution (1 M) and extracted with DCM. Compound **3** was purified using flash chromatography (0% to 40% EtOAc in hexanes) to yield a clear oil in 98% yield (510 mg, 3.42 mmol).

^1^H-NMR (400 MHz, CDCl_3_) δ 7.00 (t, J = 8.0 Hz, 1H), 6.31 (dd, J = 11.2, 8.0 Hz, 2H), 3.82 (s, 3H), 3.57 (t, J = 8.7 Hz, 2H), 2.99 (t, J = 8.5 Hz, 2H)

#### (*S*)-2-((*tert*-butyldimethylsilyl)oxy)-1-(4-methoxyindolin-1-yl)propan-1-one (compound 4)

Compound **2** (552 mg, 2.7 mmol) in DCM (1 mL) was added dropwise to a solution of compound **3** (310 mg, 2.1 mmol), DMAP (685 mg, 5.6 mmol), EDAC (451.6 mg, 2.9 mmol), and HATU (1107 mg, 2.9 mmol) dissolved in a minimal amount of MeCN. The resulting mixture was stirred overnight under argon. The reaction mixture was concentrated under reduced pressure, and the crude product was purified using flash chromatography (0% to 40% EtOAc in hexane) to yield compound **4** in 26% yield (181.7 mg, 0.542 mmol)

^1^H-NMR (500 MHz, CDCl_3_) δ 7.87 (d, J = 7.4 Hz, 1H), 7.18 (t, J = 8.2 Hz, 1H), 6.60 (d, J = 8.2 Hz, 1H), 4.57 (d, J = 6.2 Hz, 1H), 4.29-4.20 (m, 2H), 3.84 (s, 3H), 3.08 (t, J = 8.2 Hz, 2H), 1.46 (d, J = 6.6 Hz, 3H), 0.90 (s, 9H), 0.09 (d, J = 19.3 Hz, 6H)

^13^C-NMR (101 MHz, CDCl_3_) δ 171.6, 155.7, 144.8, 128.8, 118.5, 110.6, 106.2, 70.9, 55.3, 48.0, 25.8, 20.5, 18.2, −4.7, −4.8

HRMS (ESI) *m/z* calcd for C_18_H_29_N_2_O_3_Si+Na^+^: 358.1808 [*M*+Na]^+^; found: 358.1804.

#### (*S*)-2-((*tert*-butyldimethylsilyl)oxy)-1-(4-methoxy-7-nitroindolin-1-yl)propan-1-one (compound 6)

Acetyl chloride (122.2 mg, 1.64 mmol) in MeCN (1 mL) was added slowly to a solution of compound **4** (275 mg, 0.82 mmol), and silver nitrate (264.8 mg, 1.64 mmol) in MeCN (4 mL), and stirred for 3 h under an inert atmosphere. The resulting solution was filtered, and an equal volume of sodium carbonate was added. The mixture was extracted with EtOAc three times, followed by a wash with brine. The combined organic layers were dried over Na_2_SO_4_, filtered, and concentrated under reduced pressure. The crude product was purified using flash chromatography (0% to 60% EtOAc in hexane) to yield compound **5** in 10% yield (32.6 mg, 0.08 mmol) and its regioisomer compound **6** as a yellow solid in 41% yield (129.9 mg, 0.34 mmol).

^1^H-NMR (400 MHz, CDCl_3_) δ 7.79 (d, J = 9.2 Hz, 1H), 6.65 (d, J = 9.2 Hz, 1H), 4.62 (q, J = 6.9 Hz, 1H), 4.55-4.48 (m, 1H), 4.34-4.27 (m, 1H), 3.91 (s, 3H), 3.06-3.01 (m, 2H), 1.48 (d, J = 6.9 Hz, 3H), 0.93 (s, 9H), 0.14 (s, 6H)

^13^C-NMR (101 MHz, CDCl_3_) δ 172.8, 158.8, 137.7, 135.5, 125.4, 123.1, 106.3, 77.3, 77.0, 76.7, 71.4, 55.9, 50.0, 27.0, 25.7, 21.2, 18.1, −4.9, −5.2

HRMS (ESI) *m/z* calcd for C_18_H_28_N_2_O_5_Si+Na^+^: 403.1659 [*M*+Na]^+^; found: 403.1659.

#### MNI-l-lactate (compound 7)

To a solution of compound **6** (36 mg, 81 mmol) in THF (10 mL), TBAF/AcOH (1:1, 145 µl/8 µl) solution was added dropwise and stirred at 0 °C for 20 min. The reaction mixture was further stirred at r.t. for 2 h, diluted with sat. NH_4_Cl aq. and extracted with EtOAc three times. The combined organic layers were washed with brine, dried over anhydrous Na_2_SO_4_, filtered, and concentrated under reduced pressure. MNI-l-lactate was purified by flash chromatography (0% to 80% EtOAc in hexane) in a 78% yield (19.6 mg, 0.0736 mmol).

^1^H-NMR (400 MHz, CDCl_3_) δ 7.80 (d, J = 9.2 Hz, 1H), 6.69 (d, J = 9.4 Hz, 1H), 4.62-4.55 (m, 1H), 4.32-4.26 (m, 1H), 4.12 (q, J = 9.9 Hz, 1H), 3.93 (s, 3H), 3.19-3.07 (m, 2H), 1.48 (d, J = 6.4Hz, 3H)

^13^C-NMR (101 MHz, CDCl_3_) δ 174.0, 158.9, 136.5, 135.2, 125.6, 122.6, 106.8, 77.3, 77.0, 76.7, 66.4, 56.0, 49.7, 26.7, 20.6

HRMS (ESI) *m/z* calcd for C_12_H_14_N_2_O_5_+Na^+^: 289.0794 [*M*+Na]^+^; found:289.0794.

#### l-lactate assay kit

A commercial l-lactate assay kit (Lactate Assay Kit-WST, L256, Dojindo Molecular Technologies, Inc.) was used to quantify the l-lactate uncaged from MNI-l-lac upon light illumination. Before the measurements, the dye mixture stock solution, working solution containing enzyme, and lactate standard solution were prepared according to the manufacturer’s instruction. After preparing the illuminated samples, 20 µL of the sample and 80 µL of working solution were mixed in a 96-well plate, along with the lactate standard calibration samples. The plate was incubated at 37 °C for 30 minutes, and the absorbance was recorded at 450 nm using a plate reader. The unknown l-lactate concentration in each sample was back-calculated using the calibration curve.

### Characterization of caged l-lactate

A 10 mM MNI-l-lac stock solution (DMSO) was diluted to 40 µM with lac(−) buffer (30 mM MOPS, 100 mM KCl, 1 mM CaCl_2_, pH 7.2).^[15]^ The solution was transferred into a cuvette (optical path = 1 cm), and its absorption spectrum was scanned using the UV-Vis spectrometer before UV illumination (365 nm, AS ONE, handy UV lamp SLUV-4). Afterward, it was illuminated for an additional 0.5, 2, 4, 6, 8, and 10 min, and the absorption spectrum was scanned between intervals.

### Protein purification and *in vitro* characterization

The gene encoding eLACCO1 cloned into pBAD-HisB with an N-terminus 6×His tagged was expressed in *E. coli* strain DH10B (Thermo Fisher Scientific). A single colony from freshly transformed *E. coli* was inoculated into two tubes of 5 mL of Terrific Broth (1 μL mL^−1^ ampicillin) at 37 °C overnight. The seed culture was then added to 200 mL of fresh Terrific Broth (2% Luria-Bertani Broth supplemented with additional 1.4% tryptone, 0.7% yeast extract, 54 mM K_2_PO_4_, 16 mM KH_2_PO_4_, 0.8% glycerol) with ampicillin (100 μL/100 mL of Terrific Broth) in a 500 mL flask, and incubated by shaking at 37 °C and 250 rcf for 1 h to reach the exponential growth phase (OD > 0.6). Once the OD_600_ reached 0.6, l-arabinose (200 µL/100 mL TB) was added to induce expression for another 16 h in a 25 °C shaker at 250 rcf. Bacteria were then harvested and lysed using a sonicator (QSonica) for 2 cycles of 60 s sonication with a 60 s break in between at a 50% duty cycle. After centrifugation at 12,000 rcf for 20 min, the supernatant was purified with Ni-NTA affinity agarose beads (G-Biosciences) on a column (Thermo Fisher Scientific^TM^ Pierce^TM^ Centrifuge Columns 10 mL). The eluted sample was further concentrated and desalted with an Amicon Ultra-15 Centrifugal Filter Device (Merck).

For the progressive uncaging experiment, a stock solution of caged l-lactate compound (10 mM) was diluted to a final concentration of 100 µM with l-lactate (−) buffer (30 mM MOPS, 100 mM KCl, 1 mM CaCl_2_, pH 7.2). The solution was aliquoted into 5 tubes, and each tube was illuminated with 365 nm UV for 0 min, 0.5 min, 1 min, 4 min, and 8 min, respectively. To conserve protein structure, the eLACCO1 biosensor itself was not illuminated. The fluorescence response of 5 µM eLACCO1was measured in the presence of 40 µM of caged lactate (25 μL protein solution and 20 µL caged lactate were added to 5 μL lac (−) buffer). All fluorescence measurements using a plate reader were performed using the same conditions (excitation wavelength = 460 ± 20 nm).

For lactate titration experiments, sodium lactate buffers were prepared by diluting an l-lactate (+) buffer (l-lactate (−) buffer + 100 mM l-lactate, pH 7.2) in an l-lactate (−) buffer to provide l-lactate concentrations ranging from 0 to 100 µM at 25 °C. The fluorescence intensities of 5 μL purified eLACCO1 protein in 45 μL lac (+) buffer were measured in various sodium lactate concentrations ranging from 10 nM to 100 µM. For the caged l-lactate titration, a stock solution of caged lactate solution (10 mM) was diluted with an l-lactate (−) buffer to provide caged l-lactate concentrations ranging from 10 nM to 100 µM at 25 °C. Each lactate buffer was illuminated with a 365 nm UV lamp for 10 minutes to fully uncage MNI-caged lactate. The fluorescence intensities of 5 µL purified eLACCO1 protein in 45 µL caged lactate (+) were measured in various uncaged MNI-caged lactate (with 10 min UV illumination) concentrations ranging from 10 nM to 100 µM. Fluorescence intensities were plotted against both caged l-lactate and sodium lactate concentrations separately and fitted to a modified Hill Equation (equation 1) in GraphPad Prism (equation name: log(agonist) vs. response – Variable slope) to determine the apparent *K*_d_. All fluorescence measurements using a plate reader were performed using the same conditions (emission mode, the excitation wavelength = 460 ± 20 nm).

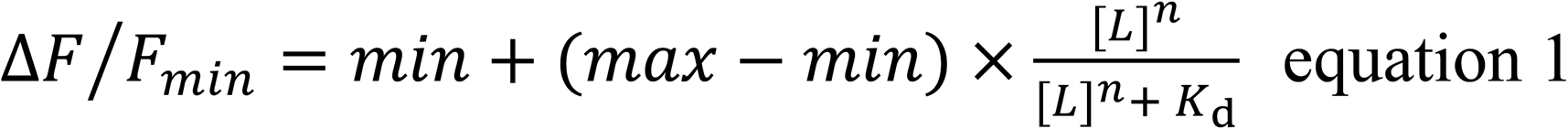

Where min and max are the minimum and maximum plateaus of the curve. [L] is the concentration of l-lactate, and *n* is the Hill slope. *K*_d_ is the dissociation constant and the lactate concentration where half the binding sites are occupied.

### Imaging of eLACCO and R-iLACCO variants with MNI-caged l-lactate in HeLa and cell lines

HeLa cells were maintained in Dulbecco’s modified Eagle medium (DMEM; Nakalai Tesque) supplemented with 10% fetal bovine serum (FBS; Sigma-Aldrich) and 1% penicillin-streptomycin (Nakalai Tesque) at 37 °C and 5% CO_2_. Cells were seeded in 35-mm glass-bottom cell-culture dishes (Iwaki) and transiently transfected with the eLACCO2.1 and deLACCO1 using polyethyleneimine (Polysciences) prior to imaging. Transfected cells were imaged using an IX83 wide-field fluorescence microscopy (Olympus) equipped with a pE-300 LED light source (CoolLED), a 40× objective lens (numerical aperture (NA) = 1.3; oil), an ImagEM X2 EM-CCD camera (Hamamatsu), Cellsens software (Olympus), and an STR stage incubator (Tokai Hit). The filter sets used in live cell imaging had the following specifications. For eLACCO variants, we used excitation 470/20 nm, dichroic mirror 490 nm dclp, and emission 518/45 nm. For R-iLACCO variants, we used excitation 545/20 nm, dichroic mirror 565-nm dclp, and emission 598/55 nm. DMD-assisted 405 nm photoactivation uncaging experiment was performed with a multi-pattern LED illumination system (Opto-line; LEOPARD2) equipped with a DMD (Mightex; Polygon 1000) and LED light source (Prizmatix; UHP-F-405 LED, 100% power). Fluorescence images were analyzed with ImageJ software (National Institutes of Health).

For the cell-based imaging of MNI-l-lac, adherent HeLa cells seeded onto a glass-bottom dish were transfected with pcDNA3.1 R-iLACCO1.2/diLACCO1 or pcDNA3.1 eLACCO2.1/deLACCO1 variants. Forty-eight hours after transfection, the eLACCO2.1/deLACCO1 imaging was performed in Hanks’ balanced salt solution (HBSS (+) Nakalai Tesque) supplemented with 10 mM HEPES, 1 μM AR-C155858 (Tocris) with 1 mM MNI-l-lac (1% DMSO) as the imaging buffer. HeLa cells expressing R-iLACCO1.2 or R-diLACCO variants were incubated with DMEM containing no glucose (Nacalai Tesque) for 3 hours prior to imaging. The imaging in the absence of an MCT inhibitor was performed in HBSS (+) no glucose solution supplemented with 10 mM HEPES and 5 mM MNI-l-lac (1% DMSO, 0.04% Pluronic F127). In contrast, imaging in the presence of MCT inhibitor was performed in HBSS (+) no glucose solution supplemented with 10 mM HEPES, and 1 μM AR-C155858 (Tocris) with 5 mM MNI-l-lac (1% DMSO, 0.04% Pluronic F127).

## Statistics and reproducibility

All data are expressed as mean ± SD or mean ± SEM, as specified in figure legends. Sample sizes (*n*) are listed with each experiment. Statistical analysis was performed using one-way analysis of variance (ANOVA) using Rstudio. Microsoft Excel software was used to plot the figures.

## Data Availability Statement

The data that supports the findings of this study are available from the corresponding authors on reasonable request.

## Author Contributions

I.M. synthesized MNI-l-lac and performed *in vitro* characterization. I.M. and T.T. decided on the photocaging group and design. I.M. and K.K.T. established the synthetic strategies for MNI-l-lac. I.M., K.K.T., Y.N., and R.E.C. designed experiments for *in vitro* and cell-based characterization and analyzed data. I.M. performed cell-based imaging under supervision from Y.K. and Y.N.. I.M., K.K.T., Y.N., T.T., and R.E.C. wrote the manuscript. T.T. and R.E.C. supervised research.

## Acknowledgments

This research was supported by the Japan Society for the Promotion of Science (JSPS) (21K14738 and 23H04151 to Y.N., 21H00273 and 23H02101 to T.T., and 19H05633 and 22H04743 to R.E.C.), and the Japan Science and Technology Agency (JST) PRESTO program (JPMJPR22E9 to Y.N.). K.K.T. is supported by the Human Frontier Science Program Cross Disciplinary Fellowship (LT000333/2021-C). I.M. is supported by the Forefront Physics and Mathematics Program to Drive Transformation (FoPM), a World-leading Innovative Graduate Study (WINGS) Program of The University of Tokyo.

## Competing Interests

The authors declare no competing interest.

## Supplementary Material

### Supplementary Figures

**Supplementary Figure 1.**
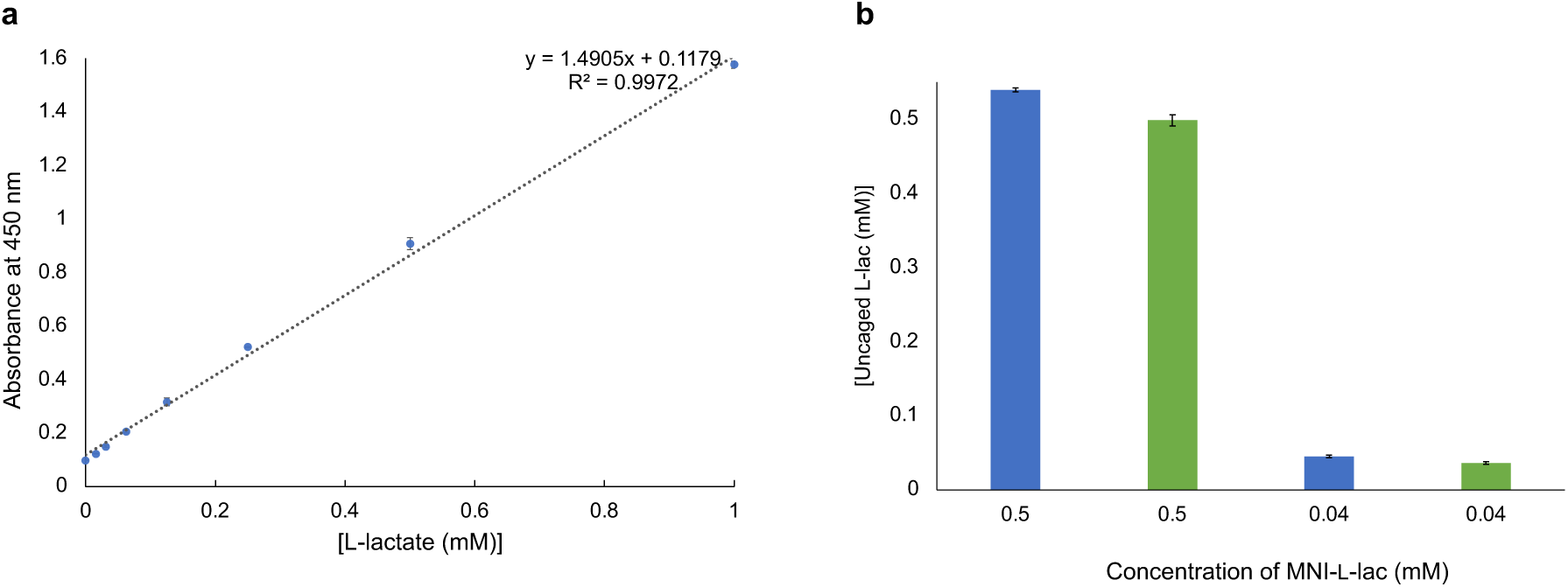
Quantification of uncaged l-lactate from MNI-l-lac using the Lactate Assay Kit-WST (L256, Dojindo Molecular Technologies, Inc.). (**a**) l-lactate calibration curve (mean ± SD, *n* = 3). (**b**) Calculated concentration of uncaged l-lactate after illumination. MNI-l-lac solution in water (blue) or lactate (−) buffer (green) was illuminated for 10 min (365 nm, 810 µW).

**Supplementary Table 1:** Released l-Lactate concentration determined using a commercial assay (mean ± SD, *n* =3 technical replicates))

| Initial [MNI-L-lac] (mM) | Uncaged L-lac in water (mM) | Uncaged L-lac in buffer (mM) |
| --- | --- | --- |
| 0.5 | $0.540 \pm 0.00269$ | $0.499 \pm 0.00740$ |
| 0.04 | $0.0448 \pm 0.00170$ | $0.036 \pm 0.00216$ |

**Supplementary Figure 2.**
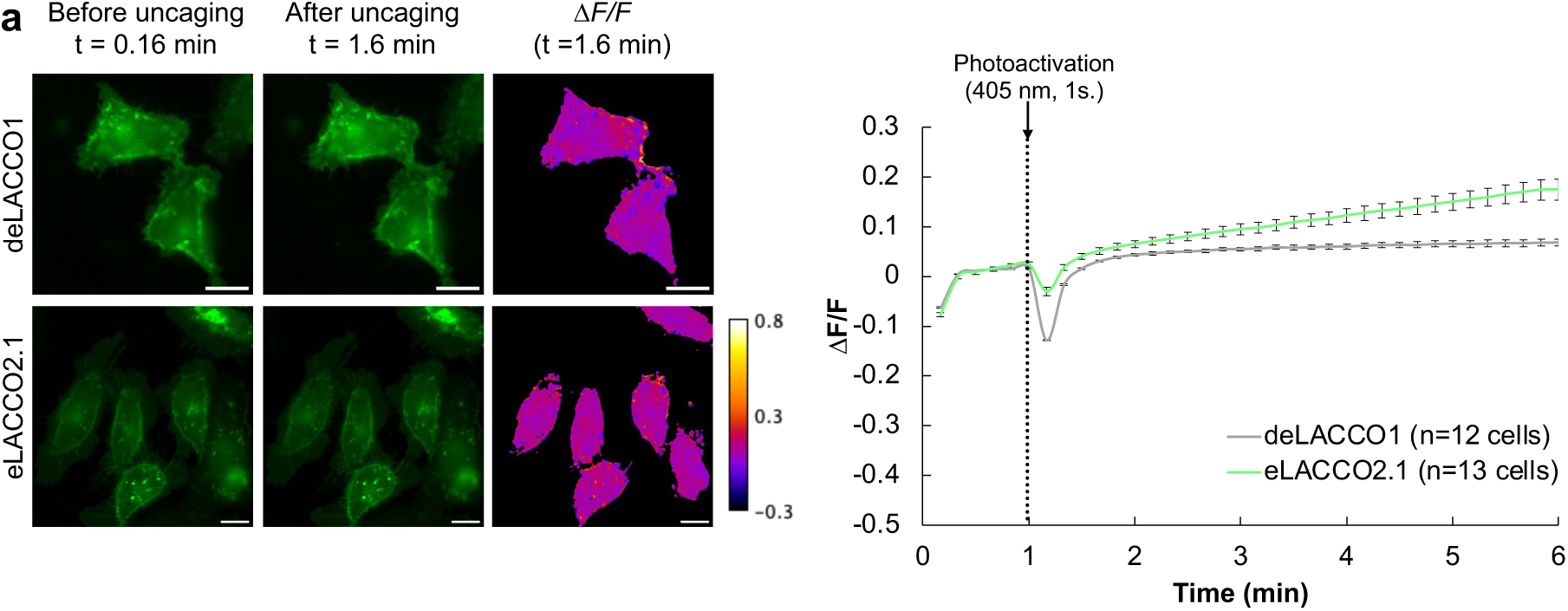
Control experiment for illumination of HeLa cells, expressing deLACCO1 or eLACCO2.1 before and after photoactivation at 1 minute in the absence of MNI-l-lac (405 nm, 1 s, 2.4 mW/cm^2^). *n* = 12 and 13 cells for deLACCO1 and eLACCO2.1, respectively (mean ± SEM). Photoactivation was performed over the entire field of view. Images in the third column are pseudocolored to demonstrate the change in fluorescence before photouncaging (*F_0_*, t = 0.16 min) and after photouncaging (*F*, t = 1.6 min). The values shown in these images were calculated as (*F-F_0_*)/*F_0_*.

**Supplementary Figure 3.**
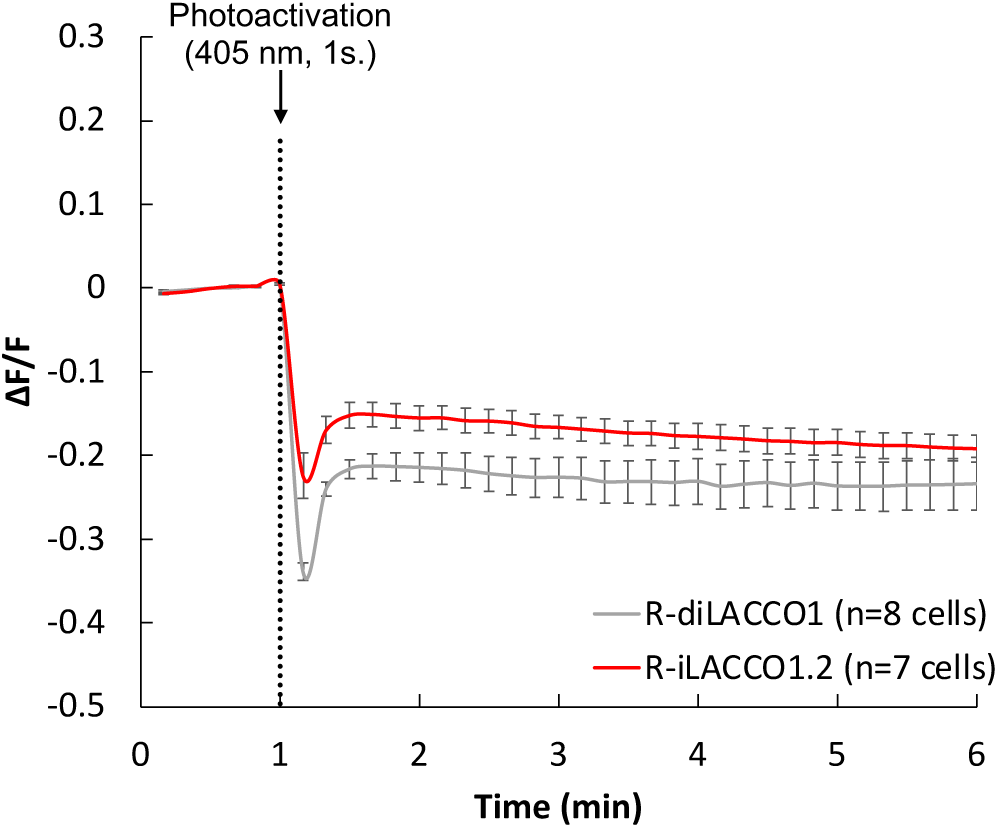
Control experiment for illumination of HeLa cells, expressing R-diLACCO1 or R-iLACCO1.2 before and after photoactivation in the presence of 5 mM MNI-l-lac and MCT inhibitor, AR-C155858. *n* = 8 and 7 for R-diLACCO1 and R-iLACCO1.2 respectively (mean ± SEM). The imaging buffer contained HBSS (+) no glucose supplemented with 10 mM HEPES and 1 μM AR-C155858 (Tocris).

**Supplementary Figure 4.**
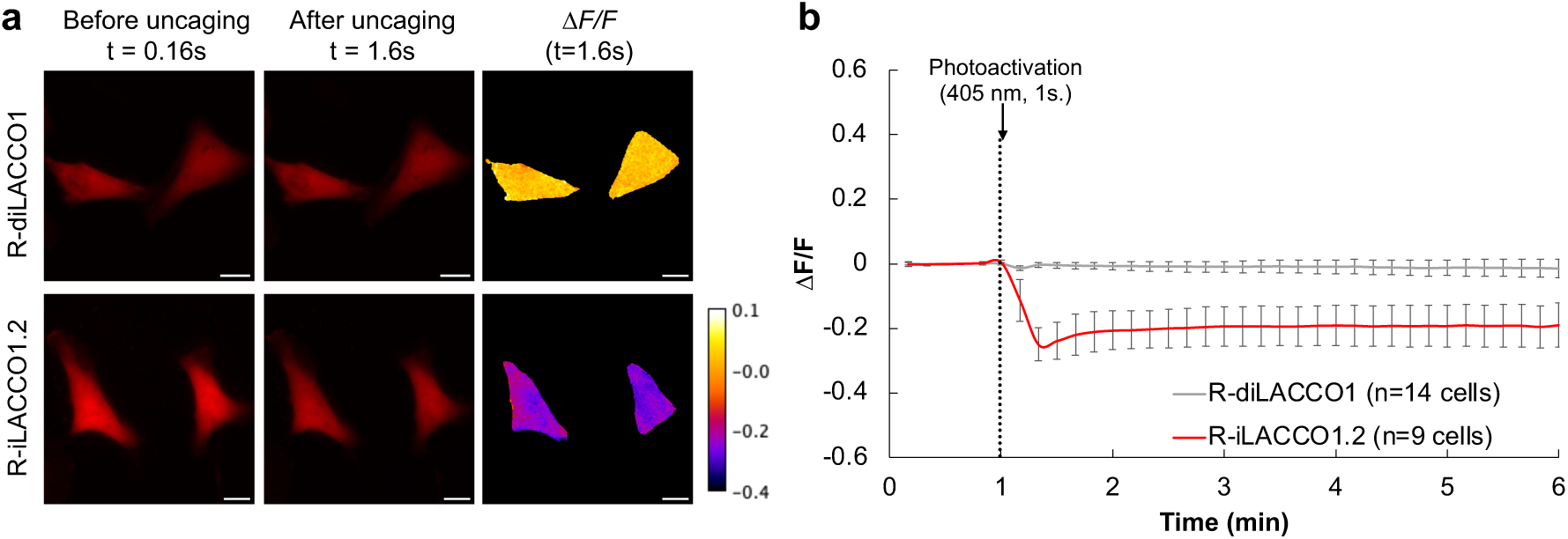
Control experiment for illumination of HeLa cells expressing R-diLACCO1 or R-iLACCO1 before and after photoactivation at 1 minute in the absence of MNI-l-lac (405 nm, 1 s, 2.4 mW/cm^2^). *n* = 14 and 9 cells for R-diLACCO1 and R-iLACCO1.2 respectively (mean ± SEM). Photoactivation was performed over the entire field of view. Images in the third column are pseudocolored to demonstrate the change in fluorescence before photouncaging (*F_0_*, t = 0.16 min) and after photouncaging (*F*, t = 1.6 min). The values shown in these images were calculated as (*F-F_0_*)/*F_0_*.

## NMR spectra for new compounds

### Compound 4 (^1^H NMR)

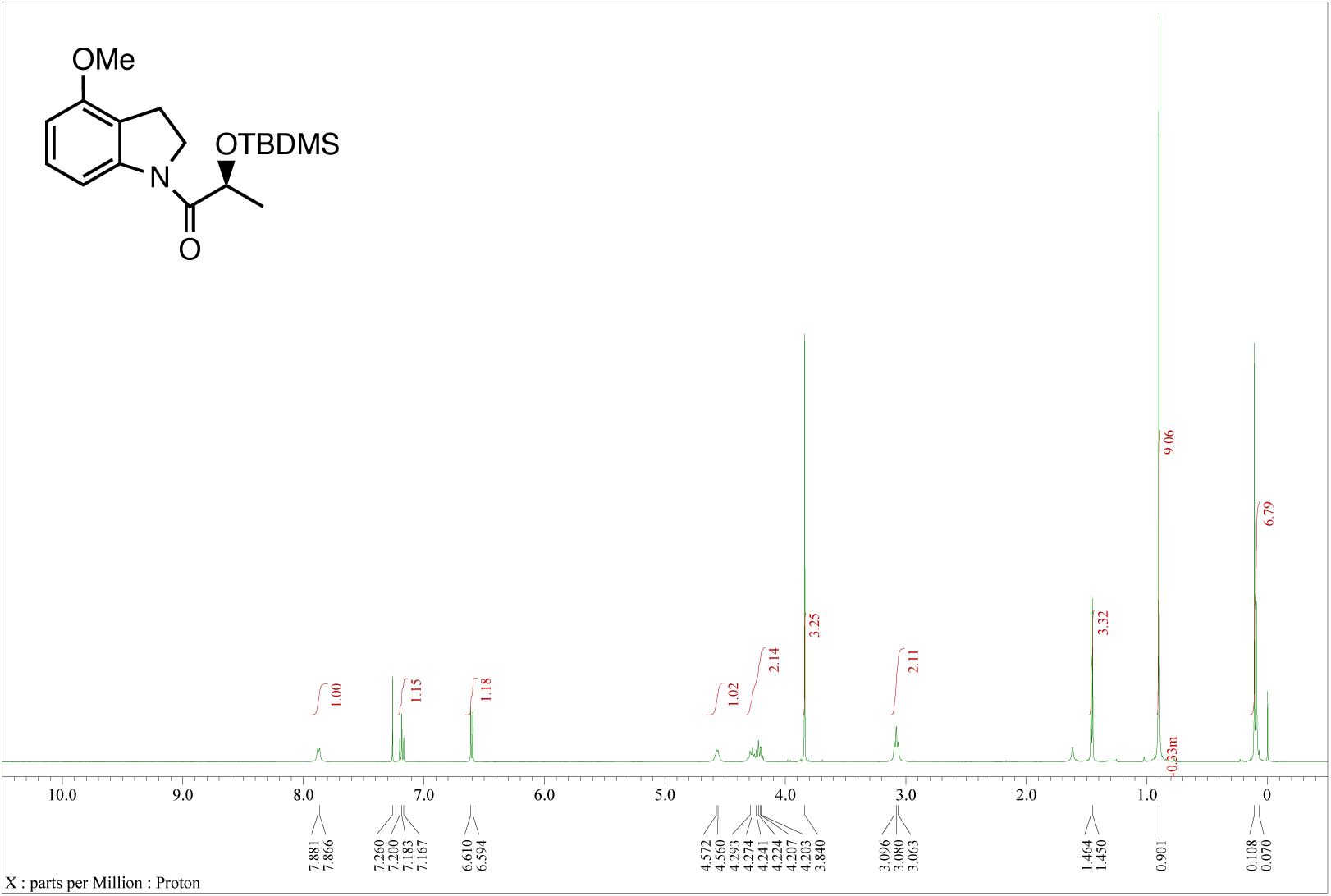

### Compound 4 (^13^C NMR)

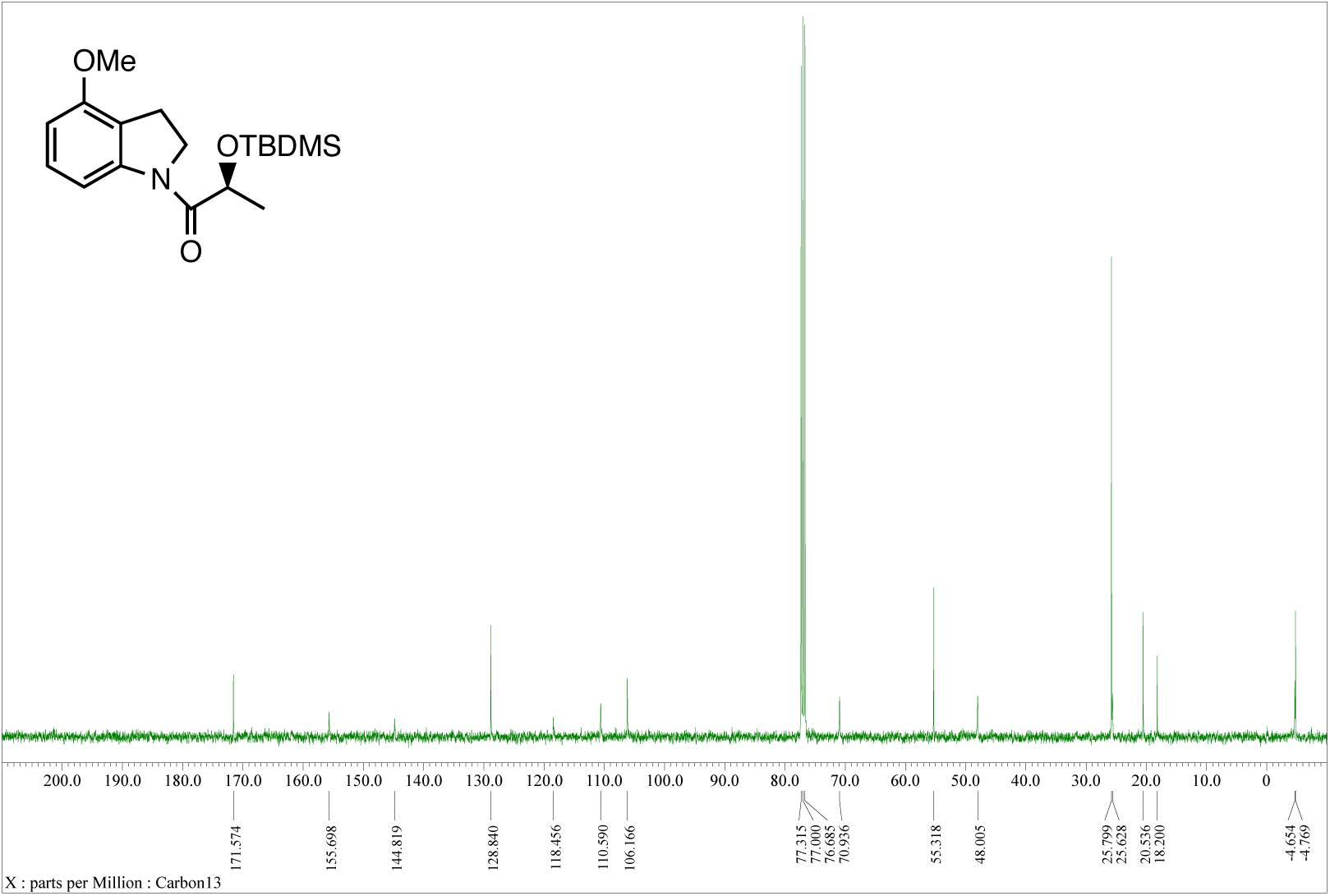

### Compound 6 (^1^H NMR)

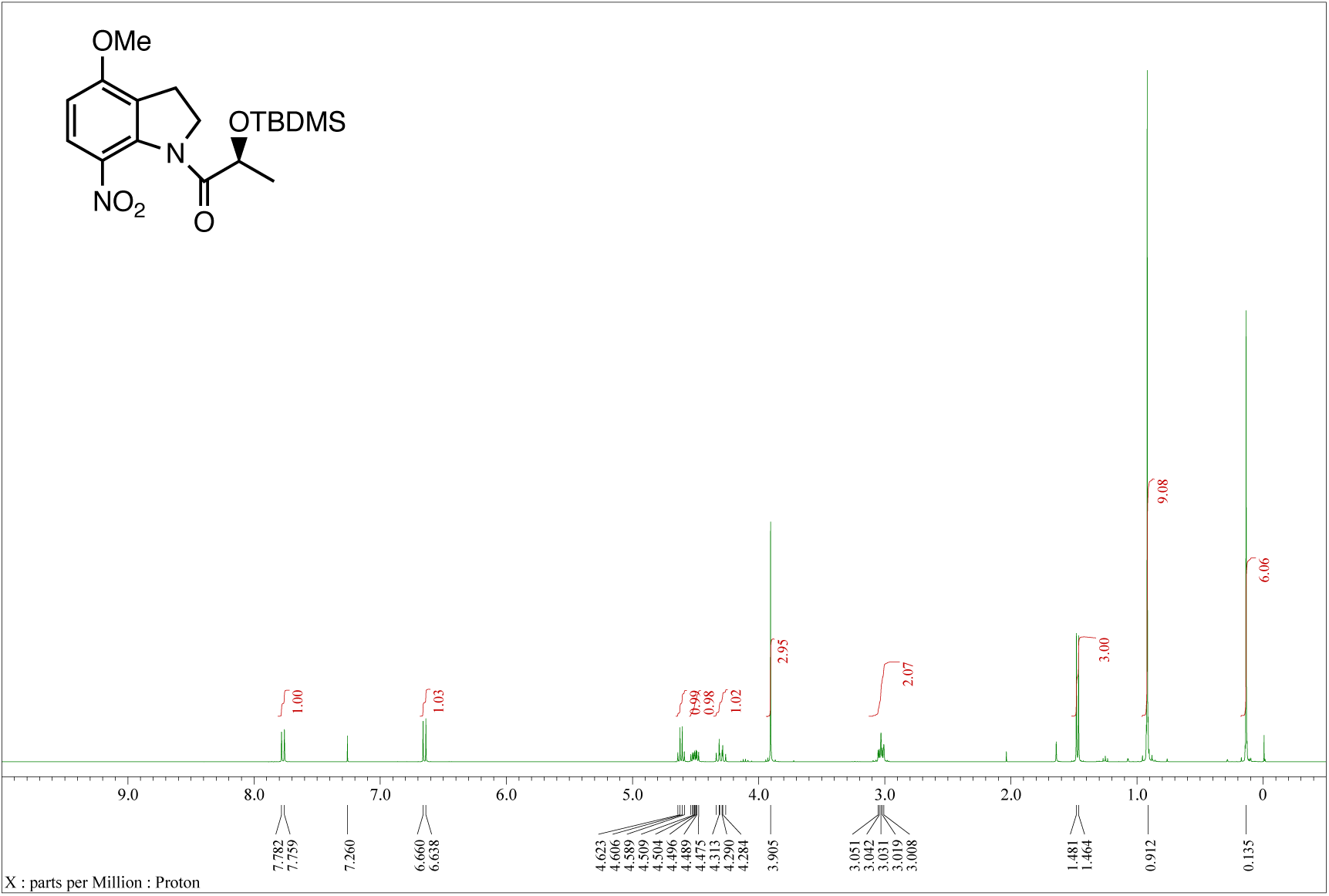

### Compound 6 (^13^C NMR)

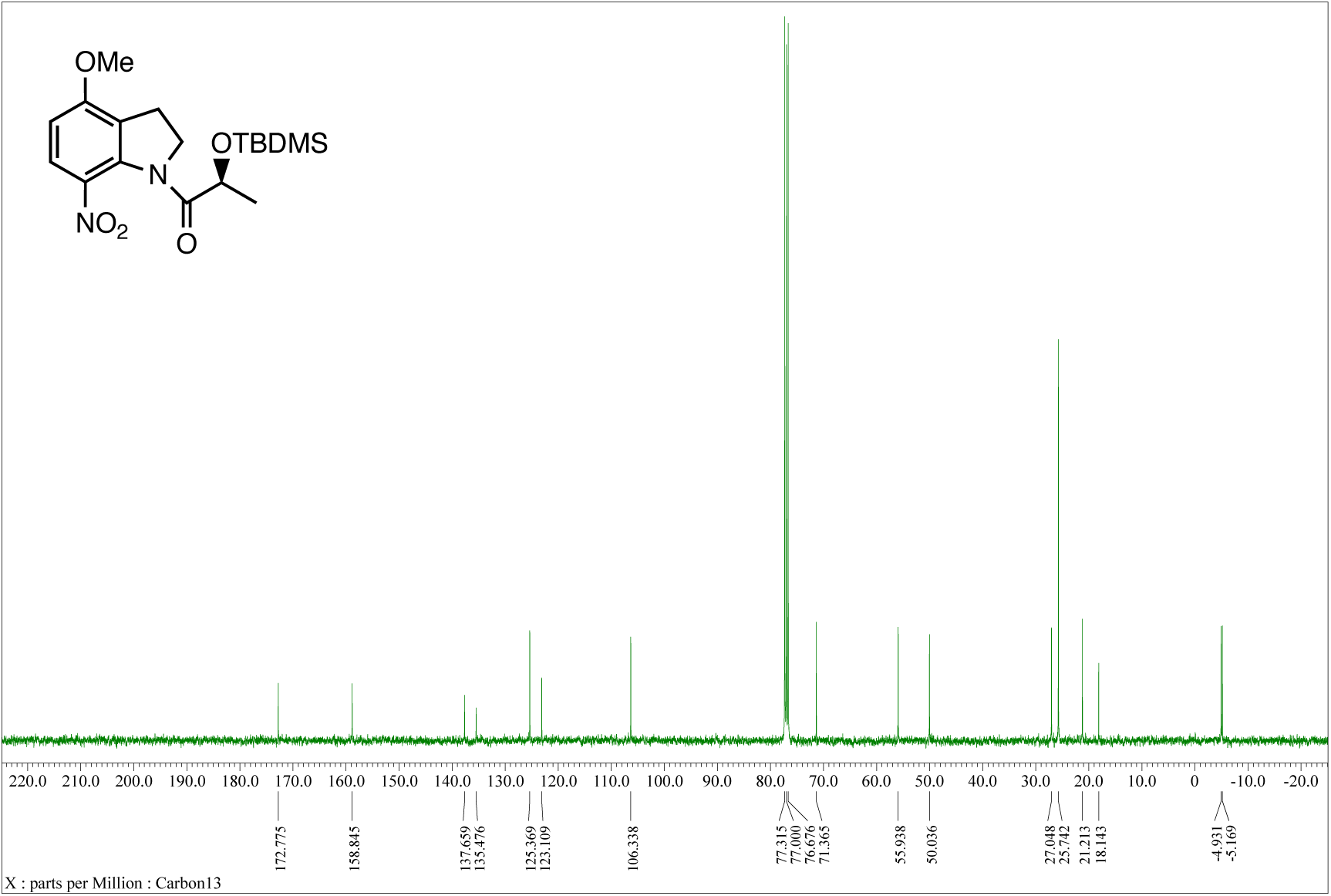

### Compound 7 (^1^H NMR)

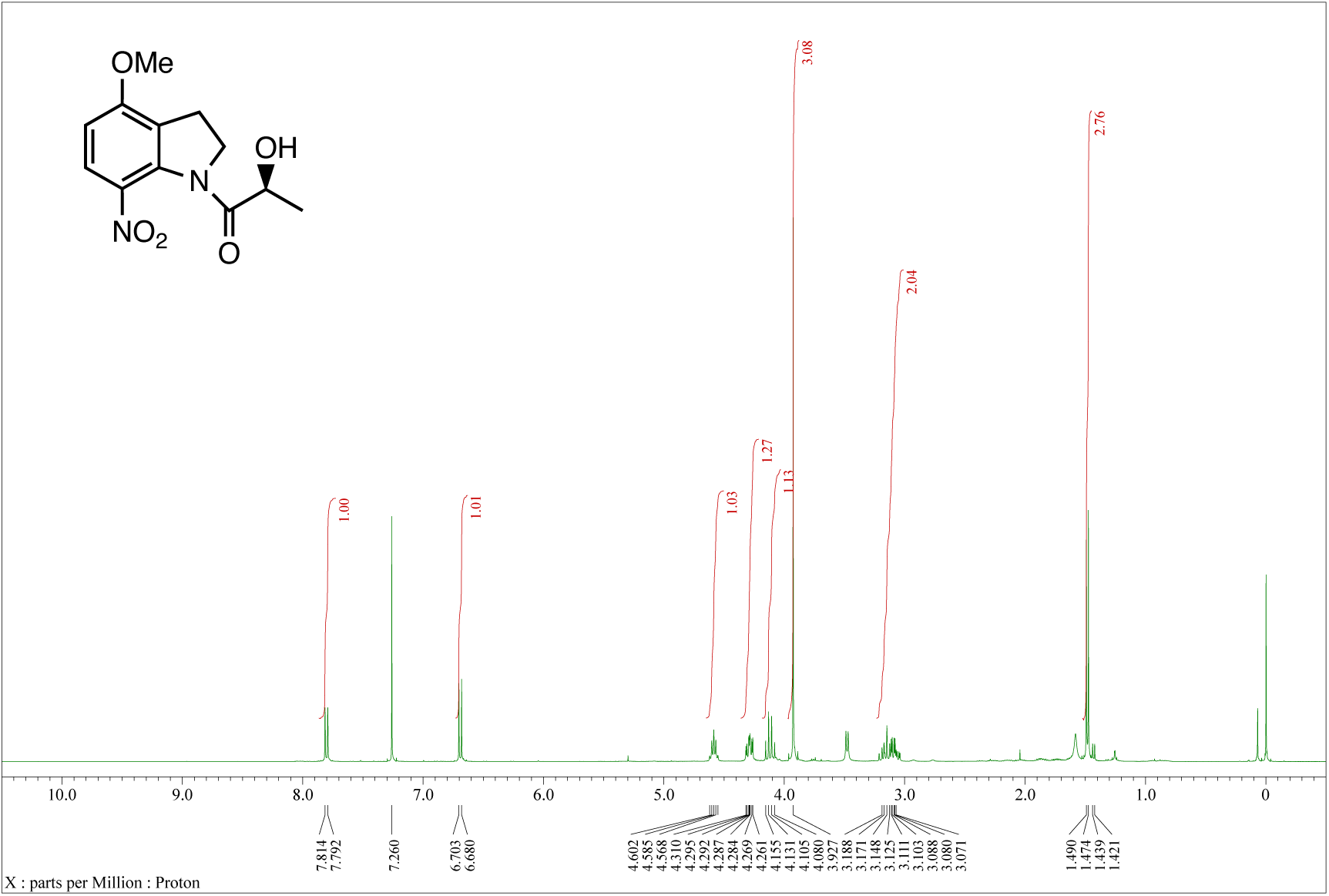

### Compound 7 (^13^C NMR)

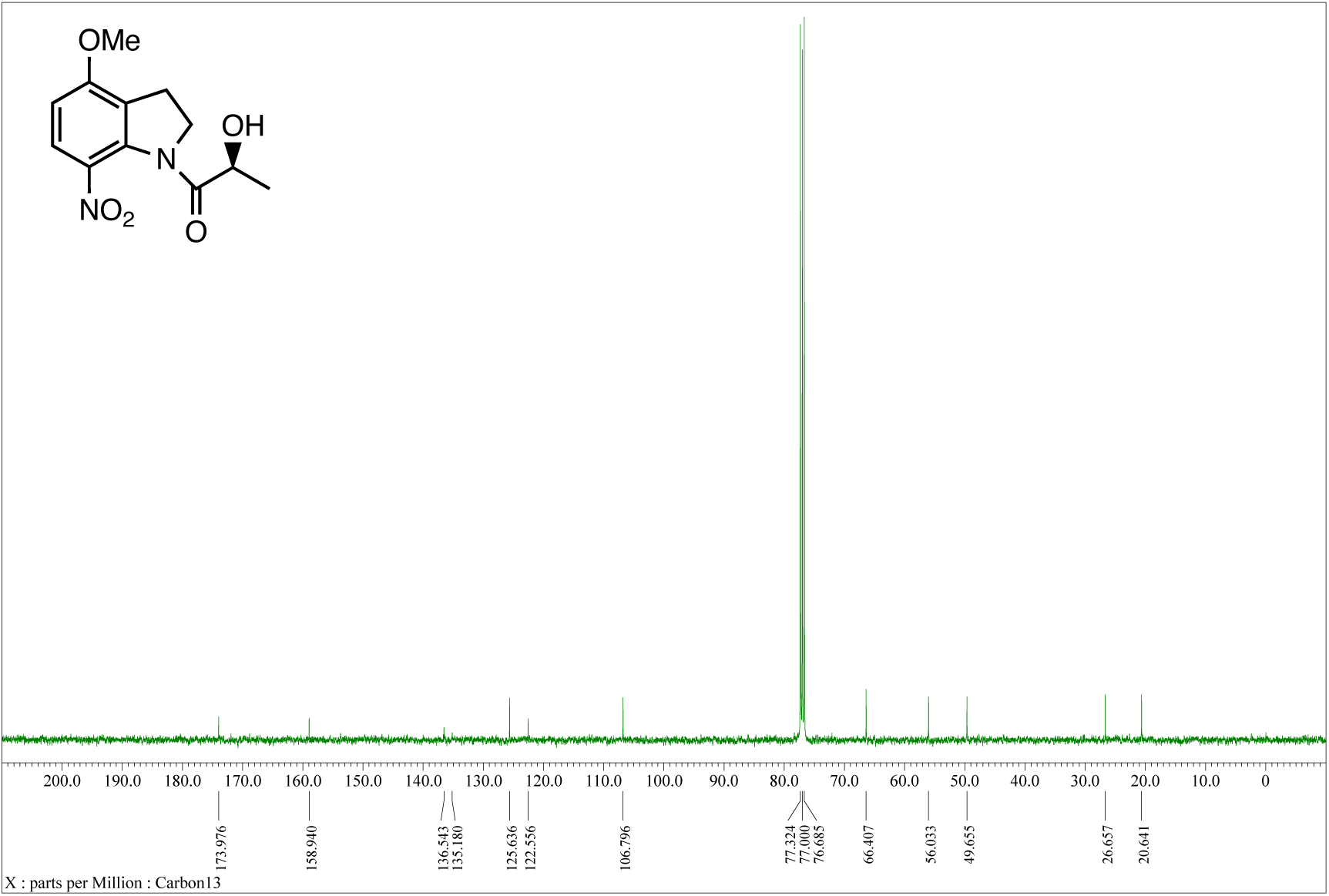

## Notes

### Competing Interest Statement

The authors have declared no competing interest.

